# Modeling Site-Specific Mutation Patterns in Pandemic-Scale Phylogenetics

**DOI:** 10.64898/2026.04.30.721865

**Authors:** Samuel Martin, Nhan Ly-Trong, Bui Quang Minh, Nick Goldman, Nicola De Maio

## Abstract

Models of genome evolution often account for different evolutionary rates at different genome positions due to, e.g., varying selective pressures or mutation rates. Recent evidence from millions of publicly shared SARS-CoV-2 genomes has revealed a more complex mutational landscape than can be modeled with existing approaches. Here, mutation rates are in fact not only highly position-specific, as currently modeled, but also nucleotide-specific; for example, specific mutations can occur very often at certain determined genome positions, while at the same positions other mutations might not be highly recurrent.

Here, we propose and investigate a general model of genome evolution where each genome position is allowed to evolve under an independent, non-normalized substitution rate matrix describing site-specific rates of all mutation types (“Site-Specific Matrix” model, or SSM). We implement SSM in the efficient pandemic-scale phylogenetic inference software CMAPLE.

Large-scale genomic epidemiological simulations suggest that, given enough data, SSM can accurately infer position- and nucleotide-specific substitution rates for more frequently observed nucleotides (typically the reference nucleotide), while other rates require higher levels of divergence. Simulations also show that SSM has a modest impact on the accuracy of phylogenetic tree estimation. We use SSM to analyze the evolution of millions of SARS-CoV-2 genomes and observe substantial mismatches between the substitution rates of classical rate variation models and our SSM estimates. These results suggest that classical models of rate variation are inadequate for modeling site-specific mutation patterns and that SSM is a useful alternative for large-scale genome analyses.

## 1 Introduction

Substitution models describe the evolution of the genetic material of a species or lineage. These models usually assume time homogeneity and are defined as continuous-time Markov processes on a phylogenetic tree. A substitution rate matrix *Q* defines such a Markov process, with each entry *q*_*x,y*_ of *Q* representing the rate at which residue *x* evolves into residue *y*. For DNA evolution models, substitution rates are embedded in a 4 *×* 4 rate matrix *Q*, representing all possible changes in the nucleotide state space {A, C, G, T}. These models, described in more detail by e.g. [Felsenstein, 2004, Yang, 2014], are central to likelihood-based phylogenetic inference methods such as IQ-TREE [Minh et al., 2020], RAxML [Stamatakis, 2014], and BEAST [Bouckaert et al., 2019].

Simple substitution models assume that a single rate matrix describes the evolution of every position (site) in a considered genomic sequence, and evolution of different sites is assumed to be independent. However, different genomic sites usually evolve at different rates due to differences in mutation rates and selective pressures [Yang, 1994b, De Maio et al., 2021]. We call this rate variation. The most popular models of rate variation assume that only the total rate of substitution *r* varies along the genome, so that the substitution rates of site *i* with total rate *r*_*i*_ are defined by the matrix *r*_*i*_*Q* [Yang, 1994b]. This means, e.g., that the relative rate of different mutation types (e.g. C to T against G to A) is constant across the genome. The rate factors *r*_*i*_ are often assumed to be independent and identically distributed according to a Γ distribution with parameters *α* and *β*. Since *β* is a scale factor, we usually set *β* = *α* so that the expected average rate across the whole sequence is 1 [Yang, 1993]. Considering many possible rates for each site can be very computationally demanding. For this reason, the gamma distribution is usually discretized with a finite number of categories *k*, frequently with *k* = 4 [Yang, 1994b]. For each site all possible *k* substitution rate matrices are then considered, leading to *k* times more calculations than a simple model without rate variation.

A more computationally efficient model of rate variation is CAT [Stamatakis, 2006] (implemented in the RAxML software [Stamatakis, 2014]), which groups sites into a number of classes, with each class assigned only one rate. This leads to a time efficiency close to a rate-homogeneous model, at the cost of ignoring uncertainty in the rate of evolution of a single site. This model is therefore better suited for alignments of many informative sequences, in which rate uncertainty for a site is low and computational demand of likelihood calculation is high. In the case that genetic data is extremely abundant, it can also make sense to avoid categorization altogether and assume a separate independent total rate parameter for each alignment position [De Maio et al., 2026].

Recent analysis of millions of SARS-CoV-2 genomes has allowed detailed investigation of the evolutionary patterns of different sites of the viral genome, and has revealed highly heterogeneous mutational [De Maio et al., 2021, De Maio et al., 2026, De Maio et al., 2025a, Haddox et al., 2025, Hensel, 2025] and selective [Bloom and Neher, 2023, Bloom et al., 2023] landscapes, with certain specific types of mutation being very recurrent at certain genome sites: for example, mutation G11083T (modifying nucleotide G into T at genome position 11083) might occur very often, but other mutations at the same site, like G11083C, might not. These patterns are a considerable source of uncertainty in phylogeny inference analyses [De Maio et al., 2025b]. This suggests that models that could account for site and nucleotide-specific rates, for example defining entirely different substitution rate matrices for different genome positions, might lead to improved evolutionary analyses. For amino acid sequences, for example, such a model has been proposed [Lartillot and Philippe, 2004] (homonymous but conceptually substantially different from the aforementioned nucleotide CAT model [Stamatakis, 2006]). This CAT model assumes that sites might evolve under one of a certain number of substitution matrices, with the number of matrices and their entries being inferred from the considered alignments.

Here, we present a nucleotide model of rate variation, “Site-Specific Matrix” (SSM), where each genomic site is assigned an independent rate matrix. This model, implemented within the highly scalable phylogenetic software package CMAPLE [Ly-Trong et al., 2024, Ly-Trong et al., 2026], combines the computational efficiency of the nucleotide CAT [Stamatakis, 2006] model with the power and flexibility of the amino acid CAT model [Lartillot and Philippe, 2004]. SSM has a large number of parameters (linear in the genome length), and is primarily meant for large informative alignments, such as SARS-CoV-2 datasets with millions of genomes. We use the SSM model to investigate an alignment of over 2 million SARS-CoV-2 sequences to construct a phylogenetic tree, estimate site- and nucleotide-specific substitution rates, and to assess impact of selective and mutational pressures on this rate rate variation.

## 2 Methods

In this section we describe the SSM model and its implementation in detail. Throughout, we will assume that the input is a set of aligned genomes, as is typical in genomic epidemiology. However, the SSM model applies just as readily to a set of genes, coding regions, or any other aligned sequences.

### 2.1 SSM Model

For each genomic site *i* (corresponding to an alignment column) we assign an independent UNREST model of nucleotide substitution [Yang, 1994a] with rate matrix *Q*^*i*^. Each non-diagonal entry 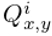 of such a rate matrix 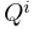 is an independent rate parameter: we do not assume that genome evolution is reversible or at equilibrium. We also do not assume that these matrices are normalized to all have the same overall mutation rate; thus different sites can naturally have different total substitution rates. We do still assume, however, that branch lengths are expressed in expected number of substitutions per site, averaged across the whole alignment. To achieve this, all *Q*^*i*^ are normalized by the same factor so that the average total substitution rate along the alignment is 1 (see Section 2.4). Finally, we define root nucleotide frequencies as equal to the genome-wide nucleotide frequencies in the reference genome.

### 2.2 Estimating Substitution Rates

We estimate the site-specific matrices *Q*^*i*^ by an expectation-maximization (EM) approach similar to [Holmes and Rubin, 2002], using the efficient short-branch approximations of [De Maio et al., 2026] applied to one site at a time. For a given site *i*, and conditional on a phylogenetic tree and starting substitution rates, each round of EM infers the expected number of mutations 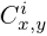 from nucleotide *x* to *y* at site *i*, and the expected waiting time 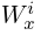 at that site for nucleotide *x* (i.e., the total expected branch length spent at nucleotide *x* for site *i*). These are inferred by traversing the tree and examining the state at site *i* assigned to each node (see [De Maio et al., 2023] for further details). Then, the new site-specific rate matrix *Q*^*i*^ is given by 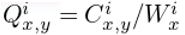 (but see Section 2.3 regarding the use of pseudocounts). These new site-specific substitution rates are used to re-infer mutation counts and waiting times at the next EM round, until convergence (defined by the log-likelihood improvement being less than 1).

As initial substitution rates for the first round of EM we use genome-wide average rates (uniform rates across sites), inferred as in [De Maio et al., 2023, Ly-Trong et al., 2024]. The procedures described in the following sections (2.3 and 2.4) are applied at each EM round.

### 2.3 Pseudocounts

In case of limited data, substitution pseudocounts [Klosterman et al., 2006] can help prevent extreme, unrealistic rate estimates, that is, reduce estimation variance. We add a pseudocount (or ‘pseudotime’) of *ω* = 1 to the waiting times of all nucleotides at all sites. This pseudocount *ω* can be adjusted in CMAPLE using the option `--waiting-time-pseudocount`. To each mutation count 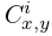 we add a pseudocount of 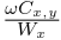, where 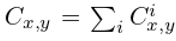 and 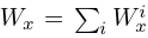 are the genome-wide total counts and waiting times respectively (and so 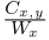 is the average observed substitution rate from *x* to *y*).

These pseudocounts prevent extreme variance in inference of substitution rates 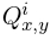, particularly when nucleotide *x* is observed only on a few branches in the tree at site *i*. In SARS-CoV-2 data, due to the low divergence of the considered genomes, this is often the case if nucleotide *x* is not the reference nucleotide at site *i*. For sites and nucleotides with very limited data, these pseudocounts will lead to inferring site-specific rates 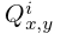 close to the genome average.

When including pseudocounts, the inferred site-specific rates become

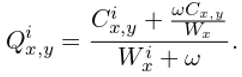

Note how 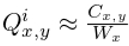 when 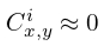 and 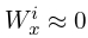.

### 2.4 Normalization

Finally, we normalize all substitution rates at all sites by one single factor to guarantee that one substitution per site is expected per unit of branch length, as is typical in likelihood-based phylogenetics, while retaining full freedom for the relative rates of all possible substitutions at all sites.

The normalization factor *r* is defined as

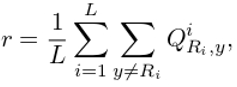

where *R*_*i*_ is the reference nucleotide at position *i*. The normalized rates 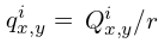 are those that we will refer to in the rest of the manuscript and that are used in CMAPLE for phylogenetic tree inference. This ensures that

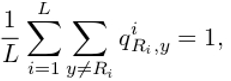

i.e., the average rate from reference nucleotides across the genome is 1, and *t* substitutions are expected per site on a short branch of length *t* descending directly from the reference genome. This normalization is intended particularly for cases in which the genomes considered have very low divergence from the reference genome, as is the case for the SARS-CoV-2 data that we consider in Sections 2.6 and 2.7.

In case of non-equilibrium and considerable divergence from the reference, the total substitution rate along parts of the tree might deviate noticeably from that of the reference genome. This means that, as with any other standard substitution model in phylogenetics, the expected number of substitutions per unit of branch length might also change along the tree.

To aid numerical stability, we bound inferred rates 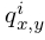 to the range 0.001– 250.

### 2.5 Implementation

The procedure described above for estimating SSM rates has been implemented in the software CMAPLE v2.0 [Ly-Trong et al., 2026] and can be enabled using the option `--site-rate MATRIX`. By default it is performed only once, after an initial tree has been constructed from the input alignment. It may also be optionally performed after each subtree prune and regraft (SPR) search of the tree that follows the initial tree construction, however we found that performing these extra rounds of estimation had little effect on the accuracy of the resulting tree (see Supplementary Figure S3). Estimated site-specific rate matrices are written to an output file and are used for phylogenetic inference (both SPR searches and branch length optimizations).

### 2.6 Simulated Data

To assess the performance of the SSM model, we simulated SARS-CoV-2-like genomes evolved under SSM using phastSim [De Maio et al., 2022]. Each genome site was simulated independently, and then all sites were collated into a single alignment. The reference genome Wuhan-Hu-1 (MN908947.3) was used as the root genome in the simulations. The simulated site-specific rates were those estimated with CMAPLE v1.2.0 on a real SARS-CoV-2 dataset (see Section 2.7).

### 2.7 Real SARS-CoV-2 Data

We ran CMAPLE on an alignment of 2,072,111 public SARS-CoV-2 genomes available in the supplementary data of [De Maio et al., 2025a]. These genomes have been consistently called with Viridian [Hunt et al., 2026] to reduce reference biases, and masked and filtered to prevent other recurrent artefacts; for a full description of this dataset see [De Maio et al., 2026].

## 3 Results

### 3.1 Simulation-based benchmark

We evaluated the performance of our SSM model on simulated SARS-CoV-2 data (see Section 2.6). We compared its performance against that of a site-homogeneous UNREST substitution model (referred to here simply as ‘UNREST’); and of a rate variation model with homogeneous relative substitution rates and one rate factor for each site (as in [De Maio et al., 2026], which we refer to as ‘rate-variation’). The SSM model, as expected, leads to a higher likelihood (Fig. 1), but the increase compared to the rate-variation model is quite limited (and not significant according to AIC and BIC) considering that the site-specific matrix model employs 12 times more free parameters (12 free parameters per site) than the rate variation model.

**Figure 1:**
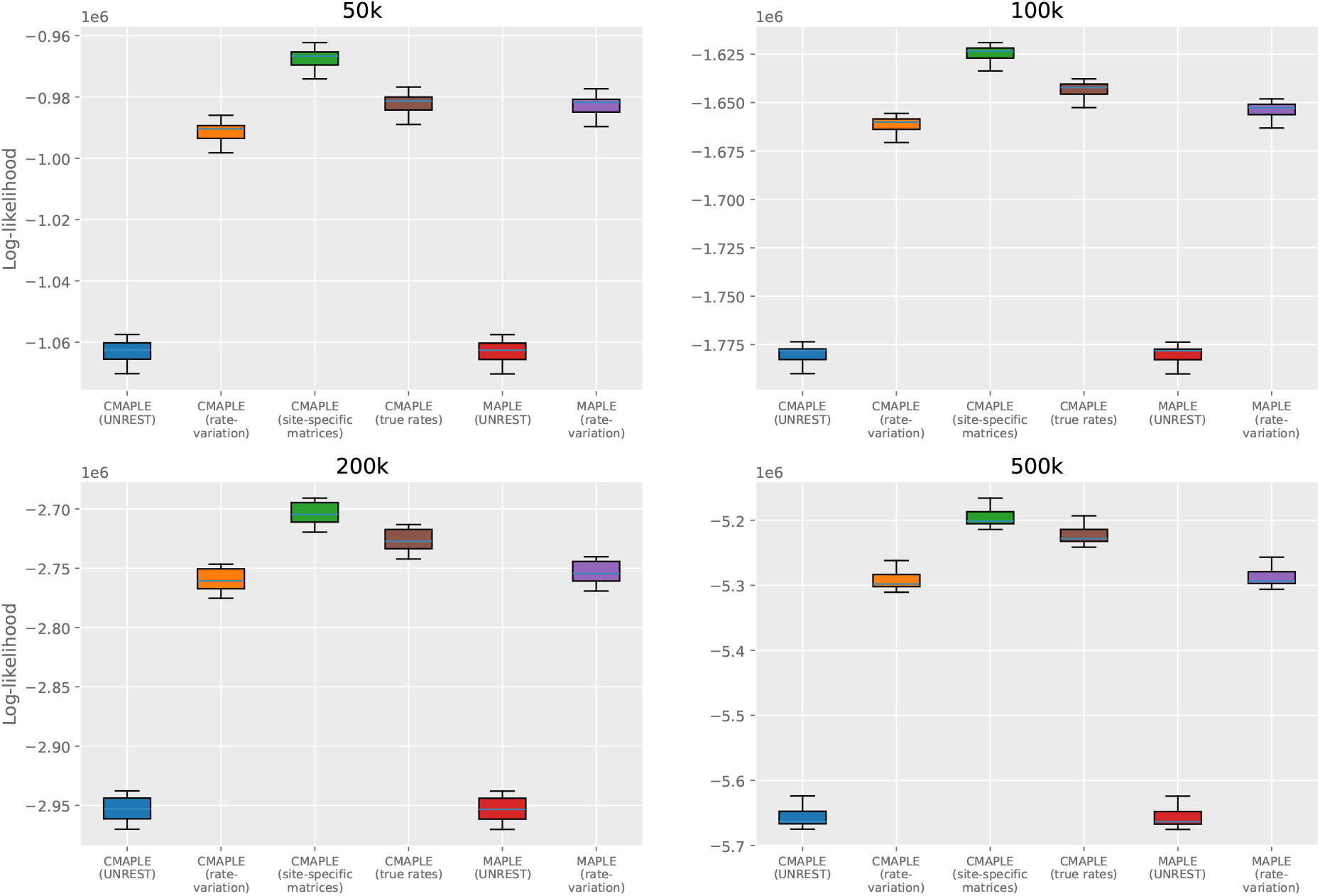
Log-likelihoods of trees reconstructed under different models. Input alignments were simulated under the SSM model from simulated trees. Ten replicate alignments were simulated for 50k, 100k, 200k, and 500k genomes respectively. For each model we optimize the substitution rate parameters, the tree topology, and the branch lengths. The models considered are UNREST (no rate variation, in red) and rate variation (in purple) as implemented in MAPLE [De Maio et al., 2023]; and UNREST (in blue), rate variation (in orange) and SSM (in green) in CMAPLE[Ly-Trong et al., 2026]. We also show in brown the log-likelihoods obtained using the correct, simulated SSM substitution rates in CMAPLE.

Trees reconstructed with inferred SSM rates are similarly accurate as those reconstructed using the correct, simulated rates (Fig. 2), suggesting negligible impact of SSM rate inference errors on phylogenetic inference accuracy. However, the increase in phylogenetic inference accuracy brought by SSM compared to other models is limited (Fig. 2). This suggests that accounting for rate matrix variation, even in ideal conditions, is not expected to substantially improve tree inference.

**Figure 2:**
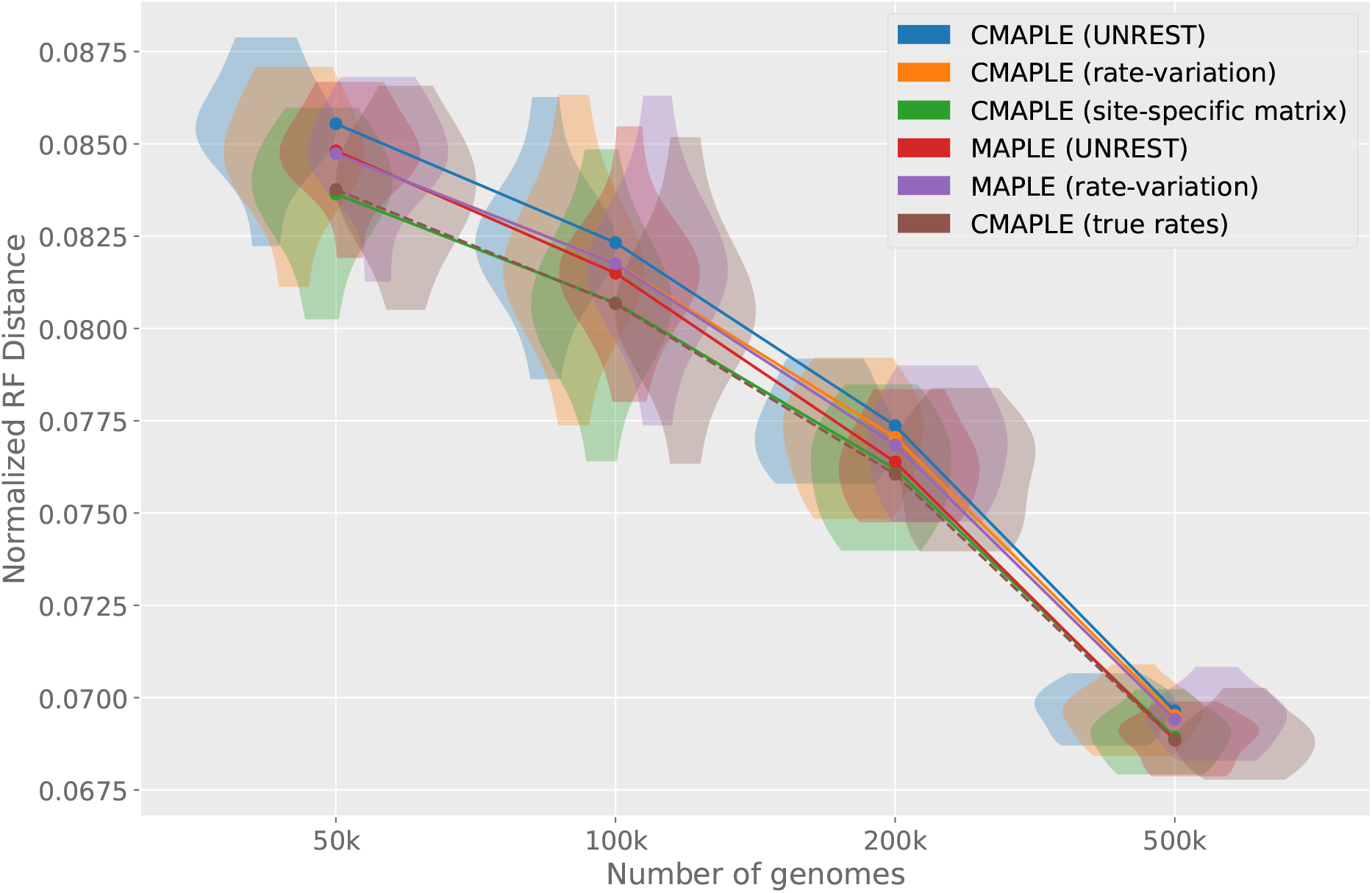
Normalized Robinson-Foulds (NRF) distances between reconstructed and true trees. Datasets and models are as in Fig. 1. Violin plots show the distribution of values across 10 replicates, whilst points show the mean values (*x*-axis values of violins are slightly offset to aid visibility). Lower NRF values mean fewer phylogenetic inference errors, and therefore more accurate phylogenetic tree reconstruction. Differences in the NRF distance between identical models in MAPLE/CMAPLE are due to features, such as error estimation, that are implemented in MAPLE but not CMAPLE.

We also evaluated SSM on data simulated under the site-homogeneous UNREST model and the ‘rate-variation’ model (see Fig. S1). As expected, the SSM model performs relatively better with simulated rate variation than without. The additional time demand for using the scalar rate variation model or the SSM model in CMAPLE, compared to a model without rate variation, is very small (see Fig. S2).

Next we evaluated the estimation of site-wise substitution rates. We split substitution rates into two groups: those from dominant nucleotides at a site, being those for which we are likely to have sufficient data to make accurate rate inference, and those from non-dominant nucleotides, for which we are less likely to make accurate rate estimations. Here, a dominant nucleotide at a particular genomic site is defined as one with waiting time of at least 25% the total tree length, meaning that the considered organism has had this nucleotide for at least a quarter of its considered evolutionary history at the considered genome position. At the short scale of divergence considered here, the reference nucleotide at the considered position is typically the only dominant nucleotide, but in some cases substitutions near the root and in branches with many descendants can result in a non-reference dominant nucleotide, or even two dominant nucleotides at the same site.

We find that estimation of SSM substitution rates from dominant nucleotides improves with more data (Fig. 3, left), and is generally more accurate than estimation of SSM rates from non-dominant nucleotides (Fig. 3, right). The rate variation model can estimate rates from dominant nucleotides reasonably well, but performs much worse with non-dominant nucleotides. The site-homogeneous model performs the worst at estimating substitution rates from dominant nucleotides, but performs better than the rate variation model at non-dominant nucleotides. The SSM model performs best with both dominant and non-dominant nucleotides (Fig. 3). Inferred rates from dominant nucleotides correlate highly with the true rates, and this correlation increases as more data is included (see Fig. 4 and Suppl. Fig. S4). Inferred rates from non-dominant nucleotides do not correlate as much, and their correlation does not improve as much as more data is considered. Since non-dominant nucleotides have typically very small waiting times, it is not surprising that rates from non-dominant nucleotides are estimated less accurately.

**Figure 3:**
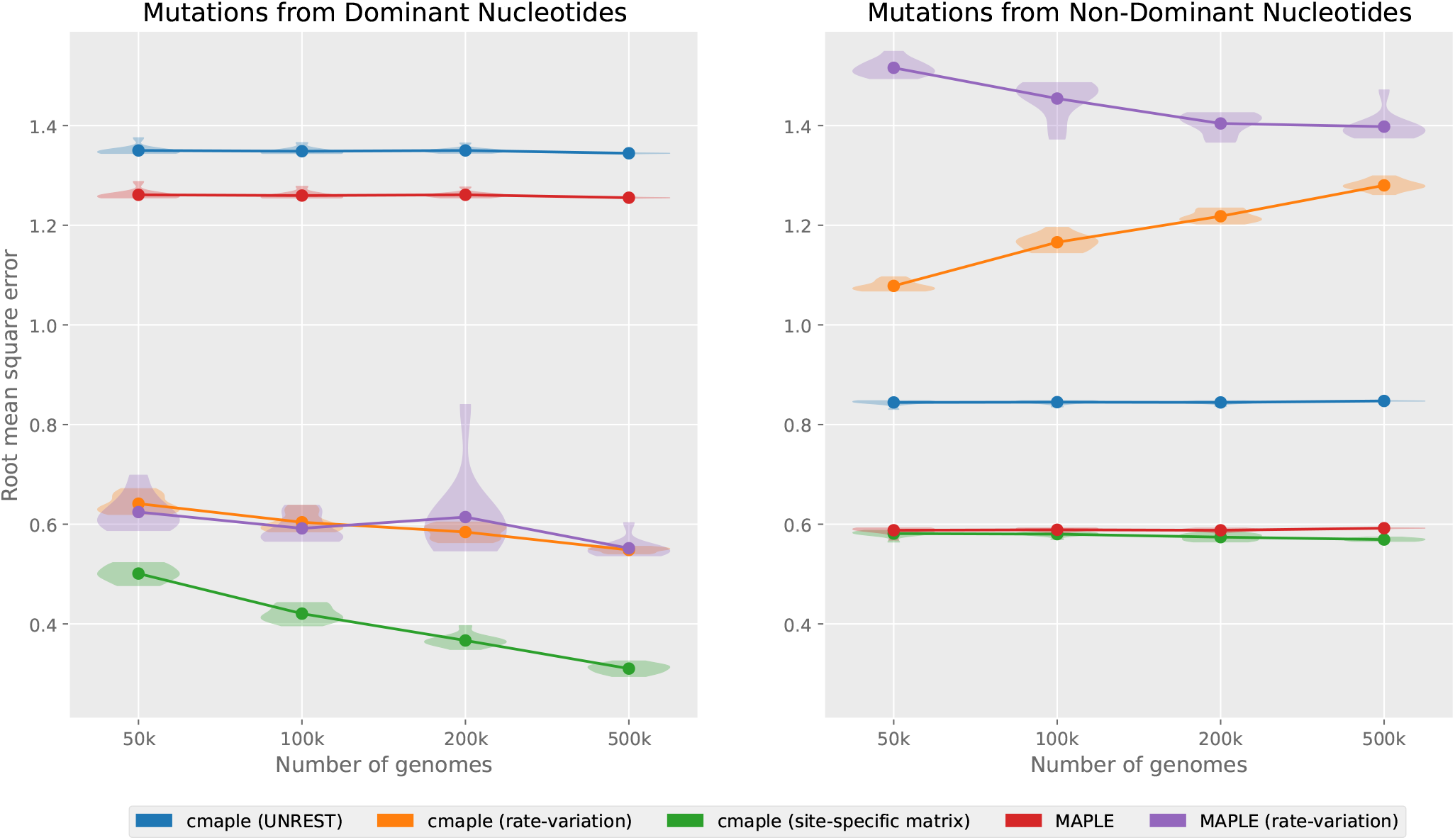
Root mean square error of all site-specific estimated rates compared to true rates, split by whether the mutation type is from a dominant nucleotide (left) or not (right) at the given site. Violin plots show the distribution over 10 replicates; points show the means. Here data was simulated under an SSM model, and the vector of inferred site-specific substitution rate matrices was compared to the vector of true simulated ones. For the non-SSM models we compare the true vector of rate matrices to the vector of site-specific rate matrices implied by the considered model (a vector of identical matrices for the site-homogeneous model, and a vector of matrices rescaled by a site-specific factor for the rate variation model). The root mean square error can be interpreted as an average difference between an estimated rate and the corresponding true rate, where the average is over all sites and rates from dominant and non-dominant nucleotides respectively.

**Figure 4:**
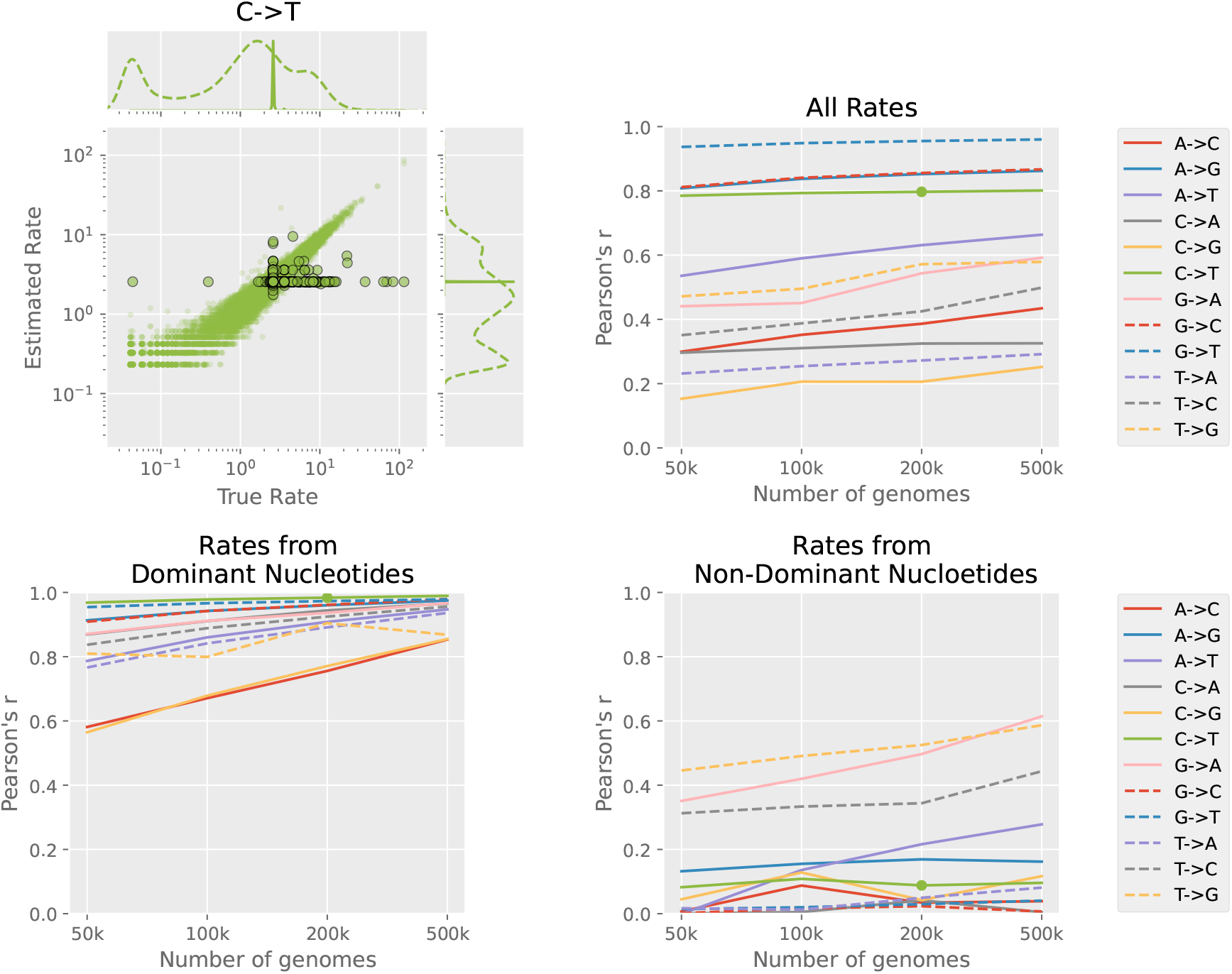
Scatter plot and Pearson’s *r* values between estimated site-specific rates and true site-specific rates. Higher Pearson’s *r* values mean more accurate substitution rate inference. **Top left** Correlation between true and inferred rates is illustrated with a scatter plot of rates of C to T mutations in a single replicate of 200k simulated data. Small circles indicate that the C nucleotide was dominant at that position, larger outlined circles indicate that it was non-dominant. Flanking plots show kernel density estimates of the distribution of true rates and estimated rates respectively, split by dominant (dashed line) and non-dominant (solid line). Note that the kernel density estimates are plotted on independent scales (not marked) and indicate only the shapes of the distributions. **Top Right** Pearson’s *r* values for all rates over all simulated datasets. **Bottom left** Rates from dominant nucleotides. **Bottom right** Rates from non-dominant nucleotides. The green point on each Pearson’s *r* plot gives the corresponding correlation from the top left scatter plot.

### 3.2 Site-specific substitution patterns in SARS-CoV-2

We compared SSM rate estimates to those from the rate variation model on a dataset of 2 million real SARS-CoV-2 genomes (described in Section 2.7). For each site *i* we have 12 rates estimated by the SSM model, one for each mutation type (denoted 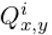 in Section 2.2). For the rate variation model, we have a scalar for each site, describing the overall mutation rate at that site. We obtain analogous mutation- and site-specific rate for this model by multiplying the site-specific scalar by the estimated genome-wide rate of the corresponding mutation type. In other words, if *Q* is the estimated genome-wide rate matrix and *r*_*i*_ is the rate at site *i*, then we define the rate of an *x* to *y* mutation at site *i* under the rate variation model to be *r*_*i*_*Q*_*x,y*_.

For each estimated SSM mutation rate 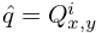, we calculated a confidence interval using a normal approximation to a Poisson distribution, where mutations are modeled as a simple Poisson process. For a given 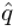, its 95% confidence interval is given by 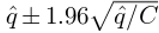, where *C* is the number of expected substitutions of the same type and at the same site along tree (see e.g. [Hogg et al., 2019, Example 4.2.2], where here 1.96 is the *z*-score for a 95% confidence interval). We consider only substitutions with at least *C* = 100 expected occurrences along the tree, of which there were 3646. For 1753 of these (48.1%) the corresponding rate (i.e., the same site and mutation type) of the rate variation model lies outside the 95% confidence interval, and for 92 of these (2.5%) the same rate is over 10 times the radius of the confidence interval away from the SSM estimate. These 92 mutations are shown in Fig. 5. (Supplementary Figure S5 shows all 1753 mutations that were outside the 95% confidence interval for comparison.) This suggests that the rate variation model is inadequate at describing highly nucleotide-specific patterns of genome sites with high mutation rates.

**Figure 5:**
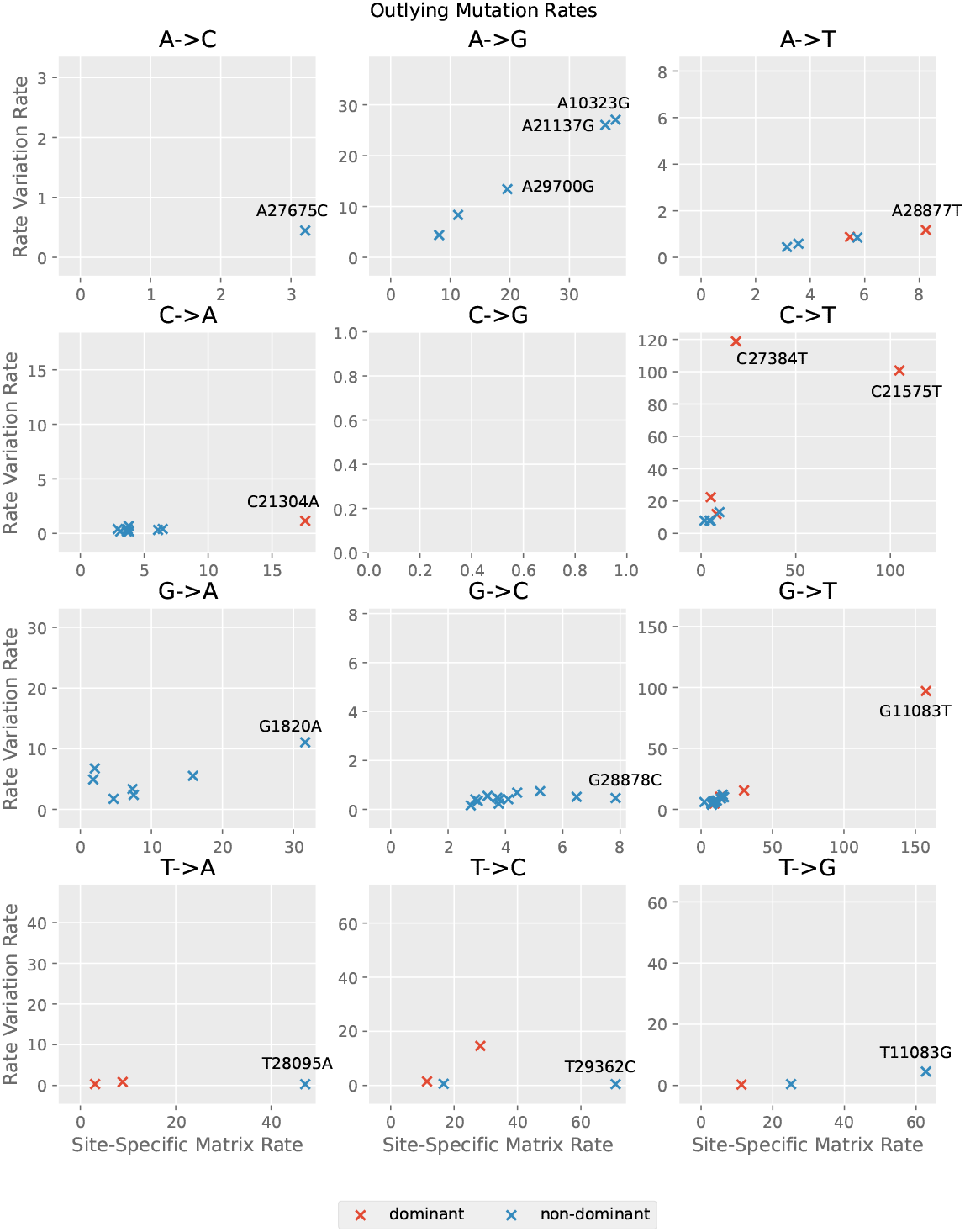
Estimated site- and nucleotide-specific substitution rates from real SARS-CoV-2 data. Estimates from the SSM model are plotted against corresponding estimates from the scalar rate variation model. Mutations from dominant nucleotides are shown in red, and mutations from non-dominant nucleotides are shown in blue. Note that for ease of visualization each subplot has a different axis range. A normal approximation was used to calculate 95% confidence intervals of the SSM estimate (not shown), and only mutations with at least 100 observations on the inferred tree were used. The plot focuses on mutations for which the scalar rate variation estimate was over 10 times the 95% confidence interval distance from the site-specific matrix estimate. We show the name of the mutations with the largest difference in inferred rate between the SSM model and the scalar rate variation model.

The mutation G11083T is generally found to be the most recurrent mutation in SARS-CoV-2 [De Maio et al., 2021], and its rate was estimated at 157.4 for the SSM model and 97.0 for the rate variation model. The rate of its reversion, T11083G, is estimated dramatically differently by the two models, at 62.8 and 4.5 respectively. This difference can be explained by the relatively low T to G background substitution rate, which is used by the rate variation model to inform the relative substitution rates at every genome site.

Similarly, SSM infers a substantially higher C21304A rate than the rate variation model (17.6 vs. 1.1). This substitution is part of a highly recurrent multinucleotide mutation [De Maio et al., 2025a], and its lower rate inferred from the rate variation model can again be explained by the low background rate of C to A mutations in general. Finally, SSM infers a substantially lower C27384T rate than the rate variation model (18.4 vs. 118.8). This mutation is the reversion of T27384C, a very frequent mutation that is also part of a highly recurrent multinucleotide mutation [De Maio et al., 2025a]. The C27384T rate in the rate variation model is likely driven up by the very recurrent T27384C mutations and by the fact that the background T to C substitution rate is relatively low.

It is natural to ask if the site-specific and nucleotide-specific substitution rate patterns are caused by variation in selective pressures or by variation in mutation rates. To address this, first we separated synonymous and non-synonymous substitutions, since non-synonymous substitutions are expected to be more strongly impacted by selection; second, we investigated the distributions of the number of descendants of substitutions, since mutations giving a fitness advantage to the virus have higher expected numbers of descendants per substitution than neutral or deleterious mutations. We found no substantial correlation between estimated substitution rates and both the mean and median number of descendants per substitution, for either synonymous nor non-synonymous mutations (Pearson’s *r* values of 0.0322 and 0.0114 respectively; see Fig. 6). We further found no substantial difference between rates at 4-fold degenerate sites and 2nd codon sites (Fig. S6). These results suggest that selective pressures have a limited impact on substitution rate variability, which is more likely impacted primarily by variation in mutation rates across sites and nucleotides. This conclusion is in agreement with [Haddox et al., 2025], who found that deviations from baseline site-specific mutation rates at SARS-CoV-2 coding sequences can be due to differences in neutral mutation rates between sites.

**Figure 6:**
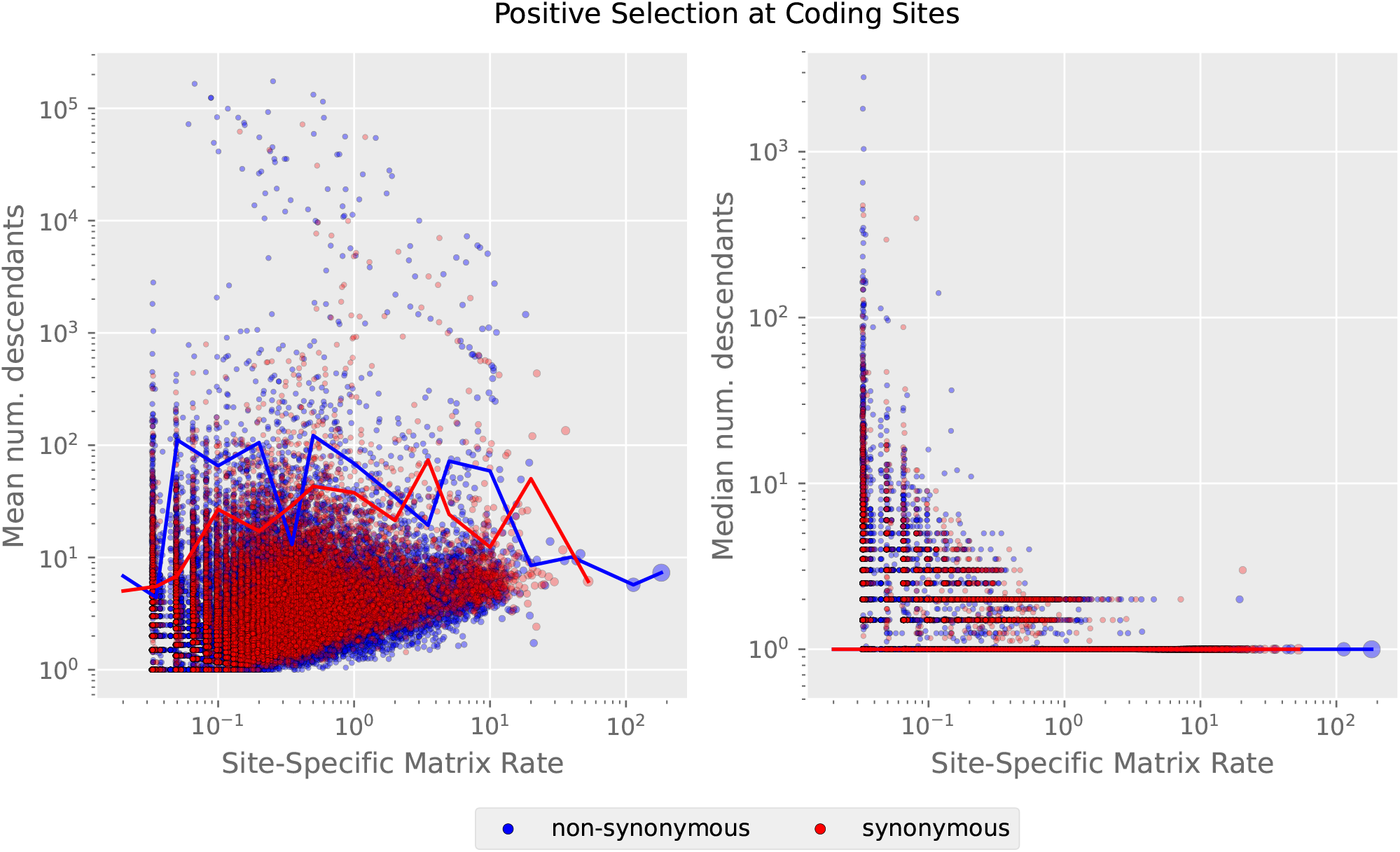
Estimated site-specific mutation rates at all SARS-CoV-2 coding sites plotted against the mean and median number of descendants for each mutation. Note the log scale for both axes. For each mutation event, the number of descendants for that event was calculated using a mutation annotated tree (MAT) output by CMAPLE, which estimates the mutations occurring on each branch of the tree under our model. We weighted the number of descendants of each mutation event by the posterior probability (between 0 and 1) assigned to the mutation event on the MAT. Mutations are coloured according to whether they are synonymous or non-synonymous, and sized by the number of occurrences on the MAT. **Left** The *y*-axis shows the mean number of descendants for each mutation type. Lines are the mean values of the number of descendants of each mutation occurring on the MAT, binned by the estimated mutation rate. **Right** The *y*-axis shows the median number of descendants for each mutation type. Lines (both at *y* = 1) are the median values of the number of descendants of each mutation occurring on the MAT, binned by the estimated mutation rate.

## 4 Discussion

We introduced SSM, a highly scalable model of nucleotide substitution rate matrix variation. This model can account for and measure strongly site-specific and nucleotide-specific mutational patterns. SSM is more heavily parameterized than other models of substitution rate variation, and while this means that it can accommodate heterogeneous substitution patterns in great detail, it also means that it requires large genomic datasets to prevent over-fitting, although we partially address this issue with the use of pseudocounts.

In our benchmark based on simulations mimicking SARS-CoV-2 evolution, our model led to slightly improved accuracy in phylogenetic inference, and resulted in trees just as good as those inferred in the ideal scenario in which the true site-specific substitution rates were known without error. However, improvements in phylogenetic inference accuracy were limited. This suggests that although mutation rates in SARS-CoV-2 are very site- and nucleotide-specific, modeling this complexity on its own does not necessarily lead to substantially better phylogenetic inference. Nonetheless, since the increase in CMAPLE run time from using SSM (or the scalar rate variation model) compared to no rate variation is small, there is little additional cost to using SSM and it enables exploration of mutation rates. We have previously observed that modeling site-specific error rates improves SARS-CoV-2 phylogenetic inference much more than modeling substitution rate variation alone [De Maio et al., 2026]. Following this observation, a potential future avenue of research could be the extension of our model to account for site- and nucleotide-specific recurrent sequence errors.

An advantage of our proposed model is that it allows us to use massive genome datasets to infer very detailed maps of mutation patterns and evolution. We showed with our simulations that we can accurately infer site-specific substitution rates when the waiting time of the mutating nucleotide is large enough (we termed these dominant nucleotides). In the considered case of SARS-CoV-2 genomes, which have low divergence, dominant nucleotides are typically those observed at the root, so we can usually accurately infer site-specific rates of mutations from the root nucleotide. On the other hand, information regarding substitution rates from non-dominant nucleotides (those with low waiting times, usually the non-root nucleotides in SARS-CoV-2) is typically limited, and our estimates of these rates are usually informed more by pseudocounts (and therefore genome-average substitution rates) than by site-specific patterns.

In the case of a traditional scalar rate variation model, these rates would instead be mostly informed by a combination of the genome-average substitution rates and by the site-specific scalar rate (which itself, in this case, would be informed by the observed substitutions and waiting times of the dominant nucleotides at the considered site). Which of these two assumptions (rates usually proportional to other rates at the same site, or to genome background rates) is more accurate might depend on the biology of the specific organisms being studied, for example on whether the variation in substitution rates is determined more by variation in mutation rates or in selective pressures. We expect that with higher-divergence datasets and given sufficiently many genomes, our model could accurately infer site-specific substitution rates for mutations from both root and non-root nucleotides.

As sequencing capacity continues to increase, and genomic epidemiology becomes more widespread, large genomic datasets and new algorithmic developments allow us to study and reconstruct pathogen genome evolution at ever greater accuracy and resolution. We expect complex and encompassing models like SSM to play a central part in the exploitation of large genome data for the detailed and scalable reconstruction, understanding, and prediction of pathogen evolution.

## 5 Data Availability

The methods described in this paper are implemented in CMAPLE v2.0, available at https://github.com/iqtree/cmaple. Simulated data, results, and code for producing the figures in this work are available at the Zenodo repository with DOI 10.5281/zenodo.19820361.

## 6 Acknowledgements

SM, NG, and NDM were were supported by the European Molecular Biology Laboratory (EMBL) and by UKRI MRC grant MR/Z503526/1. NLT and BQM were supported by a Chan-Zuckerberg Initiative grant for essential open-source software for science (EOSS4-0000000312).

---

**Figure S1:**
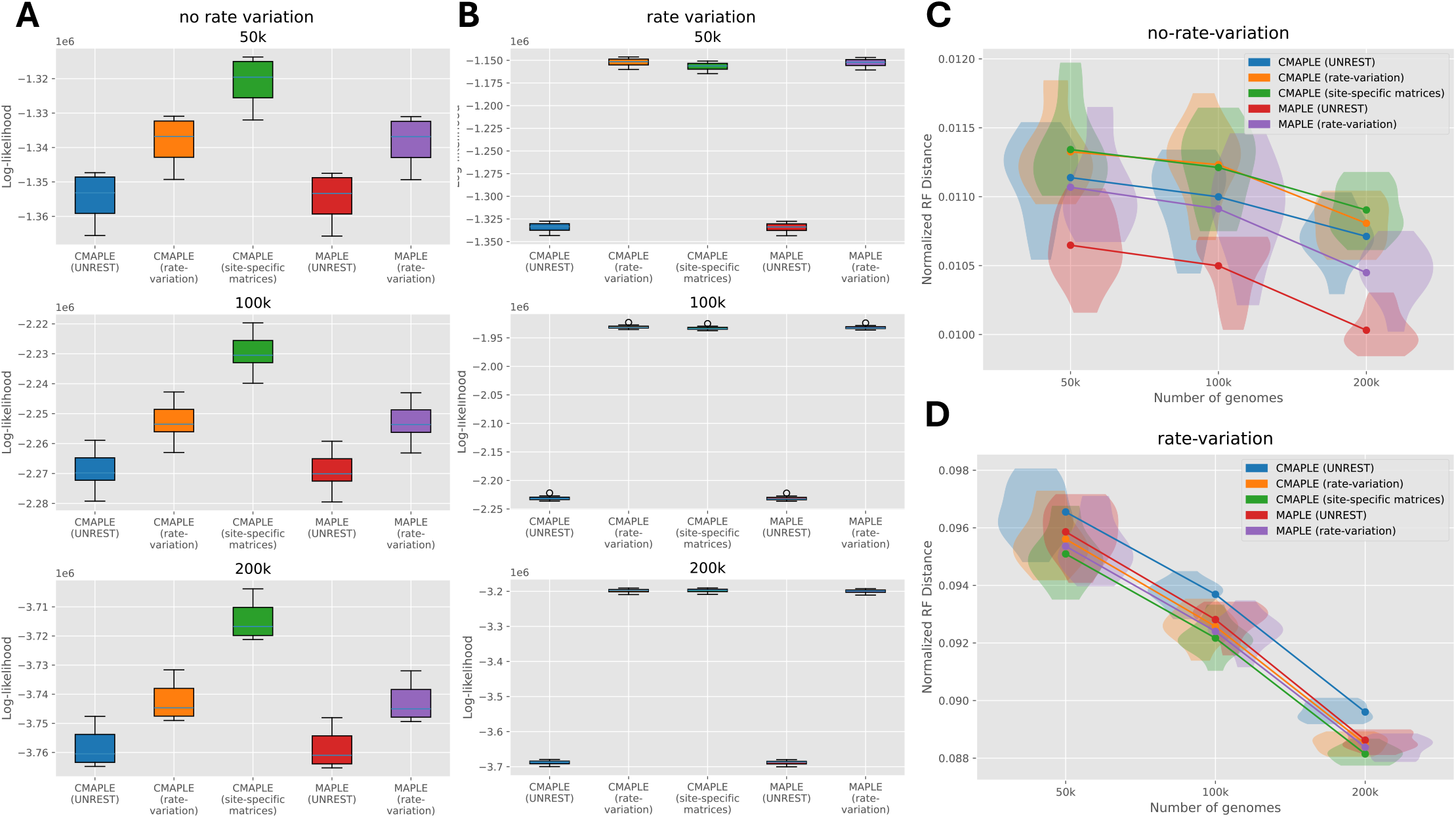
**A** Log-likelihood values of trees constructed from data simulated under the no rate variation model. **B** Log-likelihood values of trees constructed from data simulated under the scalar rate variation model. **C** Normalized Robinson-Foulds distances between the reconstructed tree and the true tree for data simulated under the no rate variation model. **D** Normalized Robinson-Foulds distances between the reconstructed tree and the true tree for data simulated under the scalar rate variation model. Alignments were simulated for 50k genomes, 100k genomes, and 200k genomes respectively. In each case we performed 10 replicates. Boxes show the interquartile range with median line. Whiskers extend to the furthest data point lying within 1.5*×* the interquartile range. Points outside the whiskers are marked individually.

**Figure S2:**
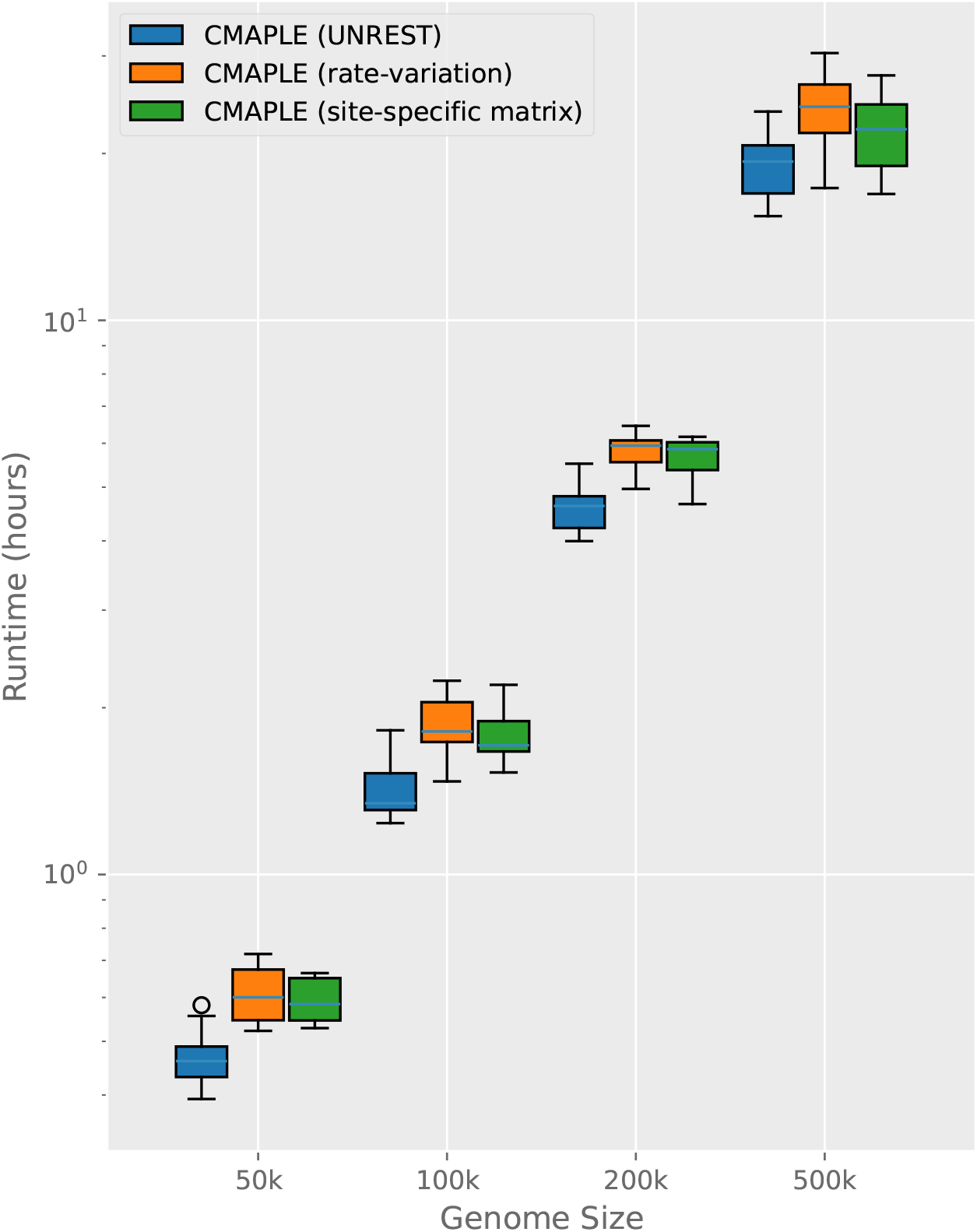
Run times of full CMAPLE analyses of simulated data for CMAPLE using a model without rate variation (blue), with scalar rate variation (orange), and SSM (green) respectively. Modelling rate variation results in only a small increase in run time. Note the log scale on the *y*-axis.

**Figure S3:**
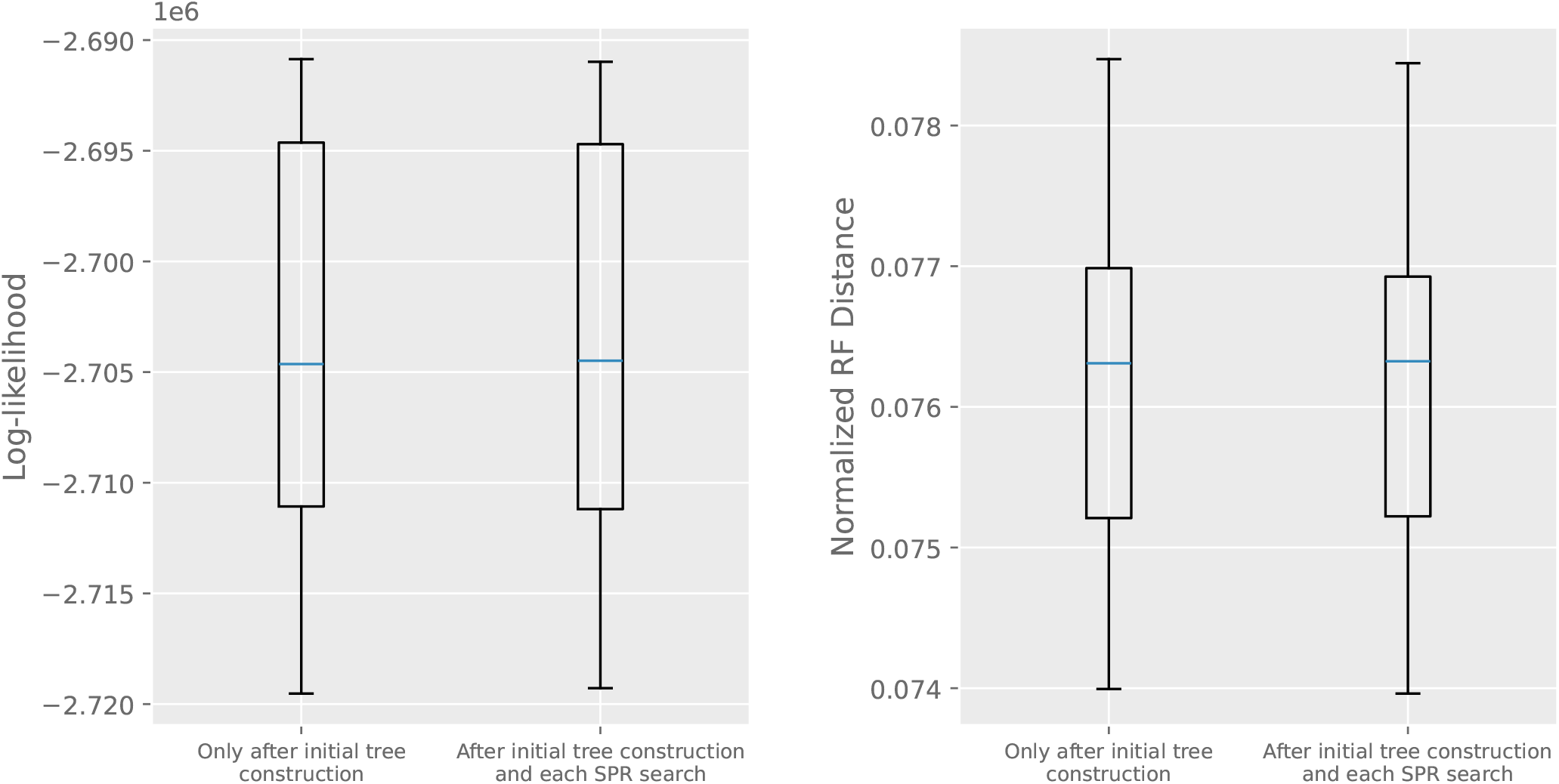
Comparison of tree log-likelihoods (left) and RF distance between the true tree and inferred tree (right) when rate estimation is performed once after the initial tree estimation, and when it is additionally performed after each subtree prune and regraft (SPR) traversal of the tree. The difference between the two is minimal. Data shown is from the 200k SSM simulated dataset. Horizontal lines in the boxes show the medians.

**Figure S4:**
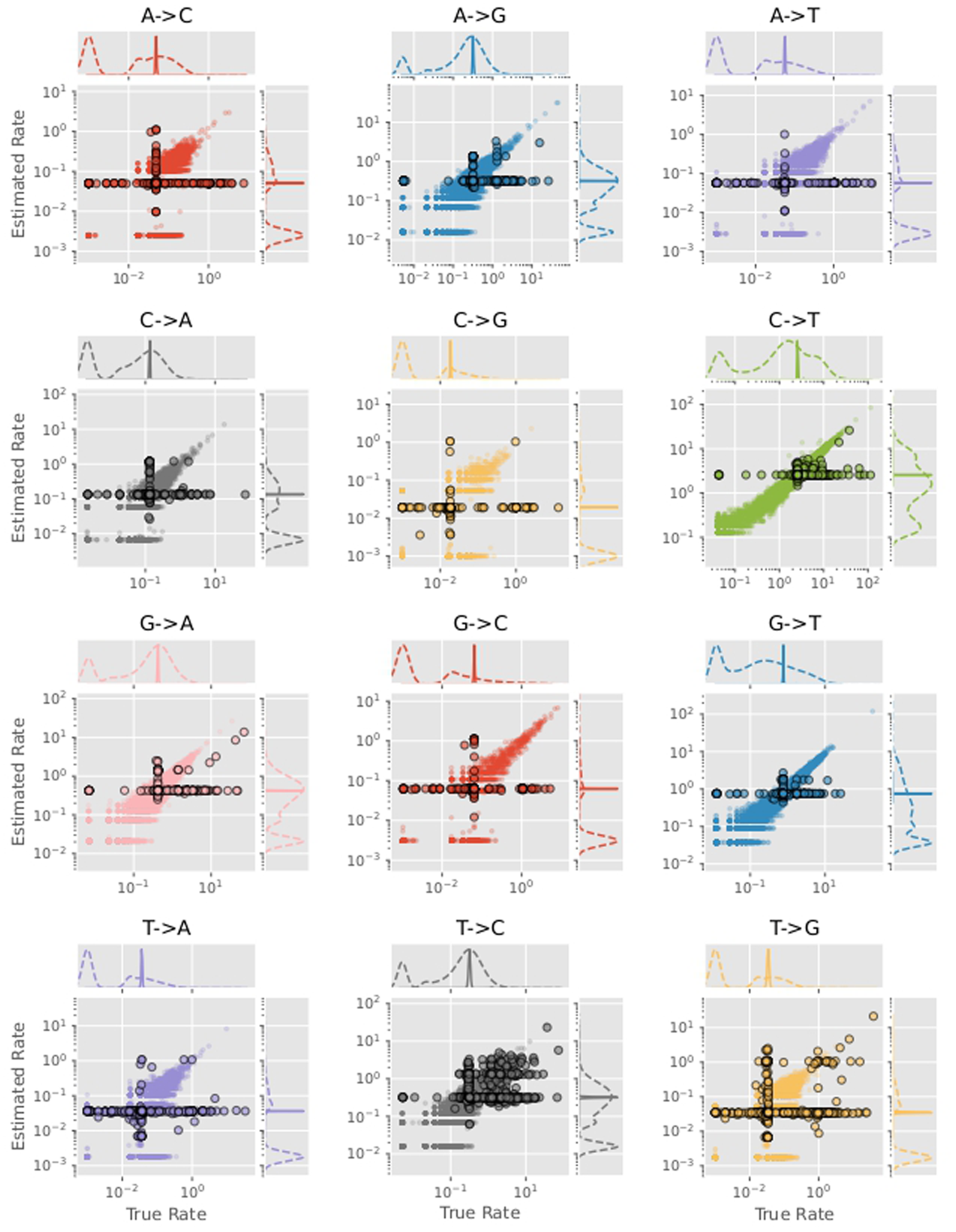
Rates from a single 500k simulated data set, split by mutation type. Each plot is further split by rates for which the mutating nucleotide is dominant at that site (marked with small circles) and non-dominant at that site (marked with larger outlined circles). Flanking plots show kernel density estimates of the distribution of true rates and estimated rates respectively, split by dominant (dashed line) and non-dominant (solid line). Note that the kernel density estimates are plotted on independent *y*-axis scales (not marked) and indicate only the shape of the distribution. Estimates of rates from non-dominant nucleotides are typically very narrowly distributed. This is because in these cases the waiting time of the mutating nucleotide is typically very small, and the estimated rates are mostly determined by the pseudocounts and are therefore close to the genome-wide average rate.

**Figure S5:**
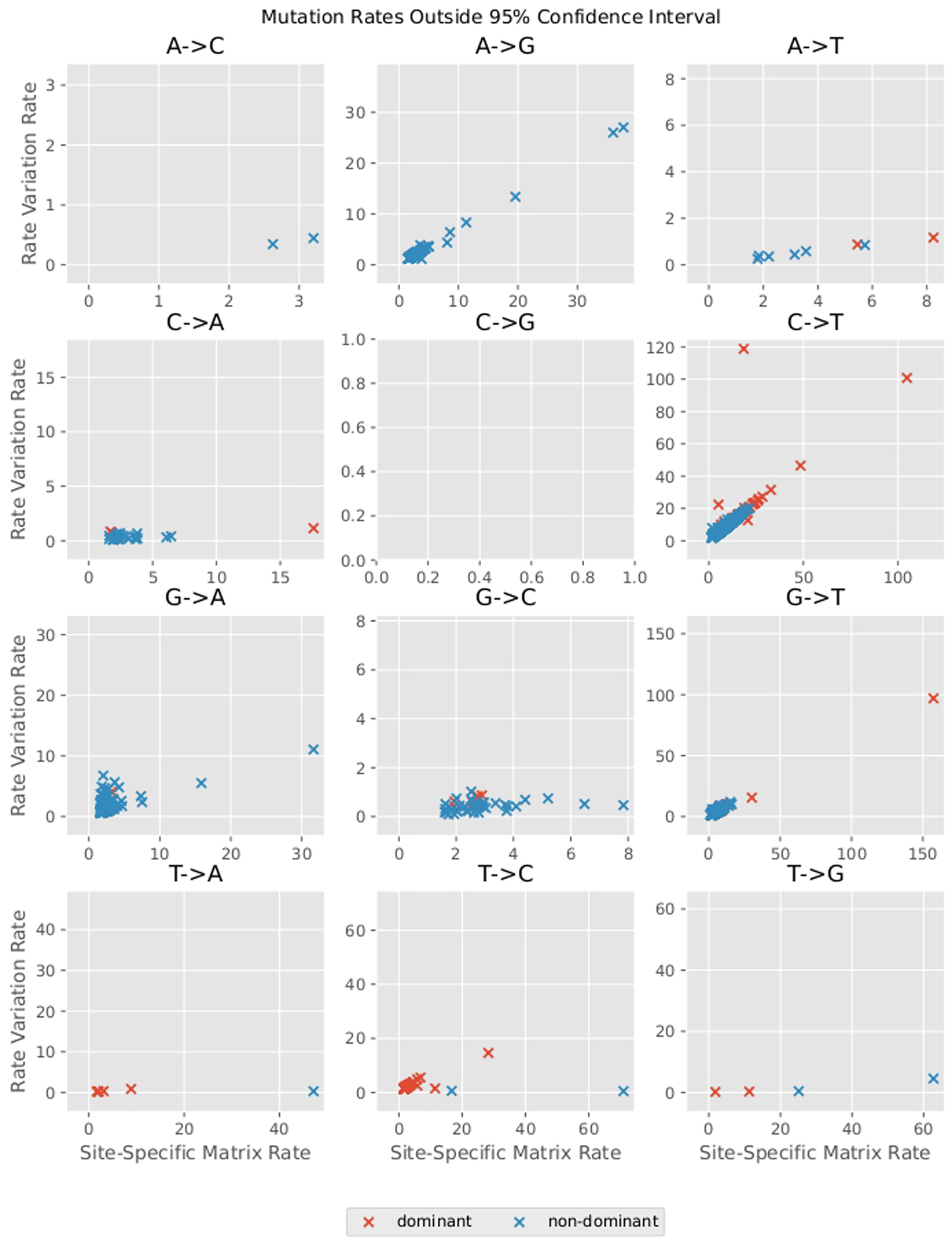
Estimated rates from SARS-CoV-2 data, as in Figure 5. Mutations from dominant nucleotides are shown in red, and mutations from non-dominant nucleotides are shown in blue. A normal approximation is used to calculate a 95% confidence interval around each SSM rate estimate. Only rates with at least 100 observations of the corresponding mutation were considered, and only rates for which the rate variation estimate is outside of the 95% confidence interval around the SSM estimate are shown (*n* = 1753).

**Figure S6:**
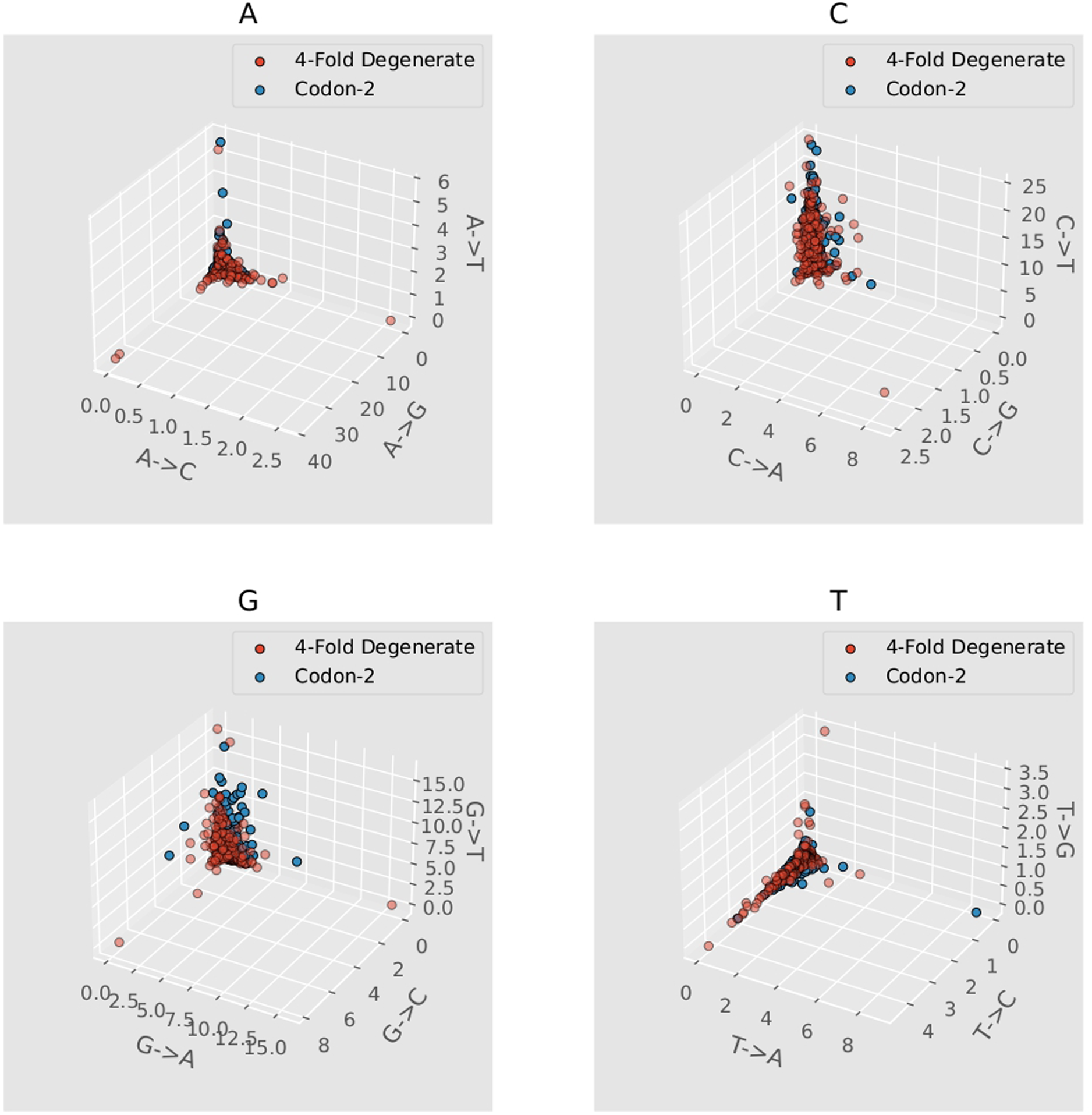
Rates at 4-fold degenerate coding sites and 2nd codon sites (“Codon-2”) in the SARS-CoV-2 genome, as estimated by SSM on the alignment of 2M real SARS-CoV-2 genomes. Each panel corresponds to a different reference nucleotide, and the three estimated mutation rates from the reference nucleotide at a given site are plotted as a 3D vector. There is no clear difference between the two distributions for any nucleotide, suggesting that variation in substitution rates is not strongly influenced by selective pressure.

## Notes

https://doi.org/10.5281/zenodo.19820360

### Competing Interest Statement

The authors have declared no competing interest.

### Summary of Updates

Added new references citing previous work estimating mutation rates in SARS-CoV-2, and updated reference for CMAPLE.

